# Frequency-specific directed interactions in the human brain network for language

**DOI:** 10.1101/108753

**Authors:** J. M. Schoffelen, A. Hultén, N. Lam, A. Marquand, J. Uddén, P. Hagoort

## Abstract

The brain’s remarkable capacity for language requires bidirectional interactions between functionally specialized brain regions. We used magnetoencephalography to investigate interregional interactions in the brain network for language, while 102 participants were reading sentences. Using Granger causality analysis, we identified inferior frontal cortex and anterior temporal regions to receive widespread input, and middle temporal regions to send widespread output. This fits well with the notion that these regions play a central role in language processing. Characterization of the functional topology of this network, using data-driven matrix factorization, which allowed for partitioning into a set of subnetworks, revealed directed connections at distinct frequencies of interaction. Connections originating from temporal regions peaked at alpha frequency, whereas connections originating from frontal and parietal regions peaked at beta frequency. These findings indicate that processing different types of linguistic information may depend on the contributions of distinct brain rhythms.

**One Sentence Summary:** Communication between language relevant areas in the brain is supported by rhythmic synchronization, where different rhythms reflect the direction of information flow.

---

The human brain is capable of effortlessly extracting meaning from sequences of written or spoken words by means of a sophisticated interplay between dedicated neocortical regions. Neuroanatomical research has revealed a number of white matter pathways that facilitate these interregional interactions (1). Electrophysiological research with electro- and magnetoencephalography (EEG/MEG) has revealed with high temporal precision the sequential activation of individual nodes embedded within the human brain network for language (2, 3). Yet, the nature of the functional interactions that enables the efficient flow of information between the nodes of this network has yet to be elucidated. Here, we show that interregional interactions in the human brain network for language are subserved by rhythmic neuronal synchronisation at specific frequencies. Specifically, we found that rhythmic activity in the alpha frequency range (8-12 Hz) propagates from temporal cortical areas to frontal cortical areas, and that beta frequency rhythmic activity (15-30 Hz) propagates in the opposite direction. These results indicate the functional relevance of rhythmic directed interactions during language processing. This functional relevance likely extends to other cognitive domains, reflecting a generic mechanism, through which information can be dynamically routed through a network of task-relevant brain regions.

One important feature of cortical interregional connections is that they are frequently reciprocal in nature (4), which implies that information can be exchanged in a bidirectional fashion. Moreover, the information flow between cortical regions may be facilitated by interregional rhythmic synchronization (5), where neuronal rhythms of specific different frequencies reflect the direction in which the information is flowing (6, 7). This bidirectional flow of information should also be a crucial feature of the neurobiological system that supports language processing. Linguistic processing is not a simple bottom-up process where incoming linguistic information (for instance, when reading a sentence) drives a sequence of activations of cortical areas that gradually transforms a string of letters into a representation of sentence and discourse meaning. Rather, contextual information, which is either already available, or built up while a sentence unfolds, can also provide top-down information, affecting the response in lower order areas.

We used magnetoencephalography (MEG) to record neuromagnetic signals while participants were reading sequences of words. We reconstructed the cortical activity in a set of predefined brain areas (consisting of 156 cortical parcels), encompassing areas that are part of the core language system, areas in the visual system, as well homolog areas in the contralateral hemisphere (Fig 1A) (8). Next, we computed frequency-resolved Granger causality to quantify directed rhythmic neuronal interactions between brain areas for language that are known to be anatomically connected (9–11). Since the interpretation of connectivity estimated from neuromagnetic recordings is highly confounded by spatial leakage of source activity (12), we statistically compared, across the sample of 102 participants, the estimated Granger causality with an estimate of Granger causality after time reversal of the signals (13). This allowed us to conservatively discard a substantial subset of the predefined connections for which the direction and/or the strength of the estimated Granger causal interaction is likely confounded by spatial leakage of activity. This left us with a subset of 713 connections from the initial 4350 connections formed between 156 modelled cortical parcels. We subsequently explored the topology of the resulting network, and observed an uneven distribution in the number of connections for the cortical parcels involved (Fig. 1B–Fig. 1C). Specifically, for each of the cortical parcels, we quantified the number of in and outgoing directed connections (i.e. the *node degree*). We observed left and right middle temporal cortical parcels to serve as a sender node in a large number of connections, projecting to ipsilateral anterior middle and superior temporal cortex (Brodmann areas (BA) 21/22/38), to contralateral middle and superior temporal cortex (BA 21/22), as well as to frontal cortex (BA 6/9/44/45/47) (p<0.05, Bonferroni corrected randomization test). Left and right inferior frontal regions (BA 47) on the other hand, were observed to receive Granger causal input from ipsilateral frontal cortex (BA 44/45/46), ipsilateral superior temporal cortex (BA 22), ipsilateral angular gyrus (BA 39), as well as ipsilateral extrastriate visual cortex (BA 19, area 17/18 present in right hemisphere only) (p<0.05, Bonferroni corrected randomization test). Additionally, regions receiving substantial inflow were located bilaterally in the anterior temporal pole (receiving input from superior and middle temporal regions, as well as from inferior frontal cortex), in the occipital pole (receiving input from extrastriate regions as well as from inferior temporal and occipito-temporal cortex), and in the right anterior temporal cortex.

**Fig. 1.**
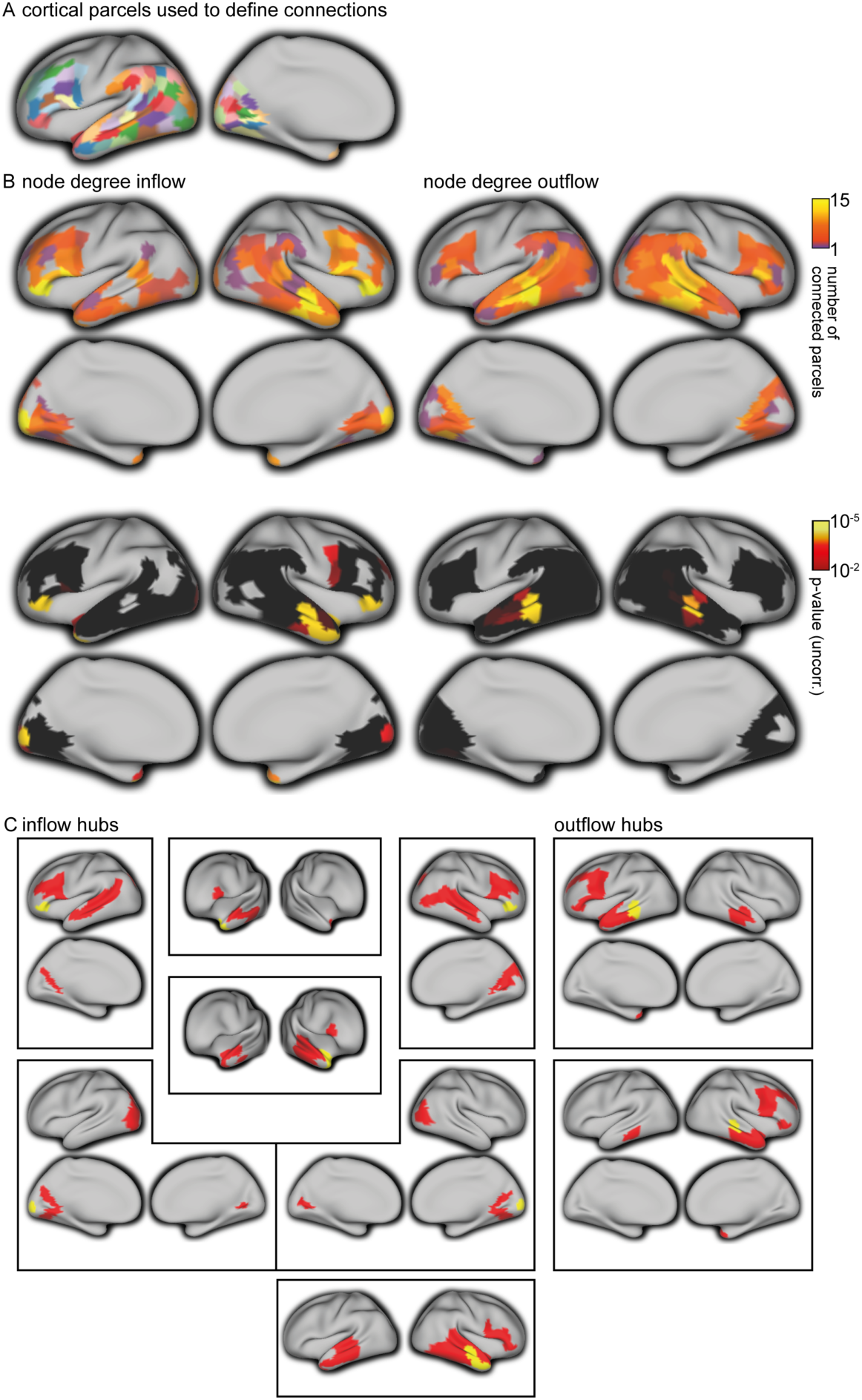
Topology of the brain network for language as quantified with Granger causality. (**A**): Overview of the left hemispheric anatomical parcels used for source reconstruction and serving as network nodes, displayed on the inflated cortical surface. Lateral and medial surfaces are shown on the left and right, respectively. Right hemispheric homologous parcels were also considered for network estimation, yet not displayed here. (**B**, upper panels): Node degree for inflow (left), i.e. the number of nodes from which each of the nodes receives significant Granger causal input (p<0.05, non-parametric permutation test, Bonferroni corrected) and outflow (right), i.e. the number of nodes to which each of the nodes sends significant Granger causal output. (**B**, lower panels): Uncorrected p-values associated with the statistical comparison (non-parametric permutation) of the topology observed in the upper panels, and randomly connected networks, keeping the overall degree distribution constant. Orange/yellow parcels survive Bonferroni correction for multiple comparisons (the number of edges), and reflect hubs in the network. (**C**): Topology of the connections for each of the highly connected hubs identified in **B**, and the other cortical areas, for inflow hubs (left panels) and outflow hubs (right panels), with the hubs displayed in yellow, and the sending/receiving areas in red.

To gain more detailed insight into the spatial and spectral structure of this brain wide network we applied non-negative matrix factorization (NMF) to the group-level connectivity data (8). Specifically, we modelled the connectivity data as a mixture of a limited number of spatially static network components, each with a subject-specific spectral profile. The decomposition algorithm did not incorporate any specific constraints with respect to the spatial or spectral structure of the underlying components. In particular, no assumptions were made about the spatial clustering of edges (i.e. the decomposition algorithm did not favor sets of connections to end up in the same component when the cortical parcels on each end of the directed connection were spatially clustered). Yet, the majority of extracted network components were physiologically interpretable, judging from the spatial clustering of the cortical parcels participating in component-specific directed interactions. Figure 2 (A-H) shows the network components with predominant connections between language-relevant cortical areas (components with predominant connections between visual cortical areas, and components with more spatially diffuse connections are shown in Supplementary figure 2). The components’ cortical locations for outflow and inflow are depicted in blue and orange/yellow, respectively, in the leftmost panel for each quadruplet of columns. For some of the components, the subject-averaged spectral profiles were band-limited to a certain frequency range, which moreover showed a consistent peak frequency across subjects (Fig. 2 B-H, middle panels for each quadruplet of columns). This suggests that these components represent frequency-specific rhythmic directed interactions between key regions in this large scale network. We categorized the extracted components based on the dominant region for outflow. The majority of the components reflected predominantly intrahemispheric connections (Fig 2 B-H, right panel for each quadruplet of columns). We identified left and right hemispheric directed rhythmic interactions from posterior and mid-temporal cortical regions to ipsilateral frontal cortex (mainly inferior frontal), with a median peak frequency at 12 Hz (interquartile range (IQR) 11-13 Hz) (Fig. 2B). A somewhat spatially more diffuse component with predominantly left intrahemispheric connections led from mid-temporal areas to inferior and superior frontal areas (Fig. 2C). Connections from posterior and mid-temporal regions to ipsilateral anterior temporal cortex had a slightly higher median peak frequency of 14 Hz (with an IQR of 12-15 Hz, and 13-15 Hz for the left and right hemispheric components), as compared to the temporo-frontal connections (Fig. 2D). Next, there was a set of components predominantly interconnecting temporal cortical regions, and which showed somewhat more variability in their spectral profile across subjects (Fig. 2E). These components reflected connections from superior and middle temporal cortex (along the whole anterior-posterior axis) to mid and anterior inferior temporal cortex, and connections from mid-middle and superior anterior temporal cortex to the temporal pole.

**Fig. 2.**
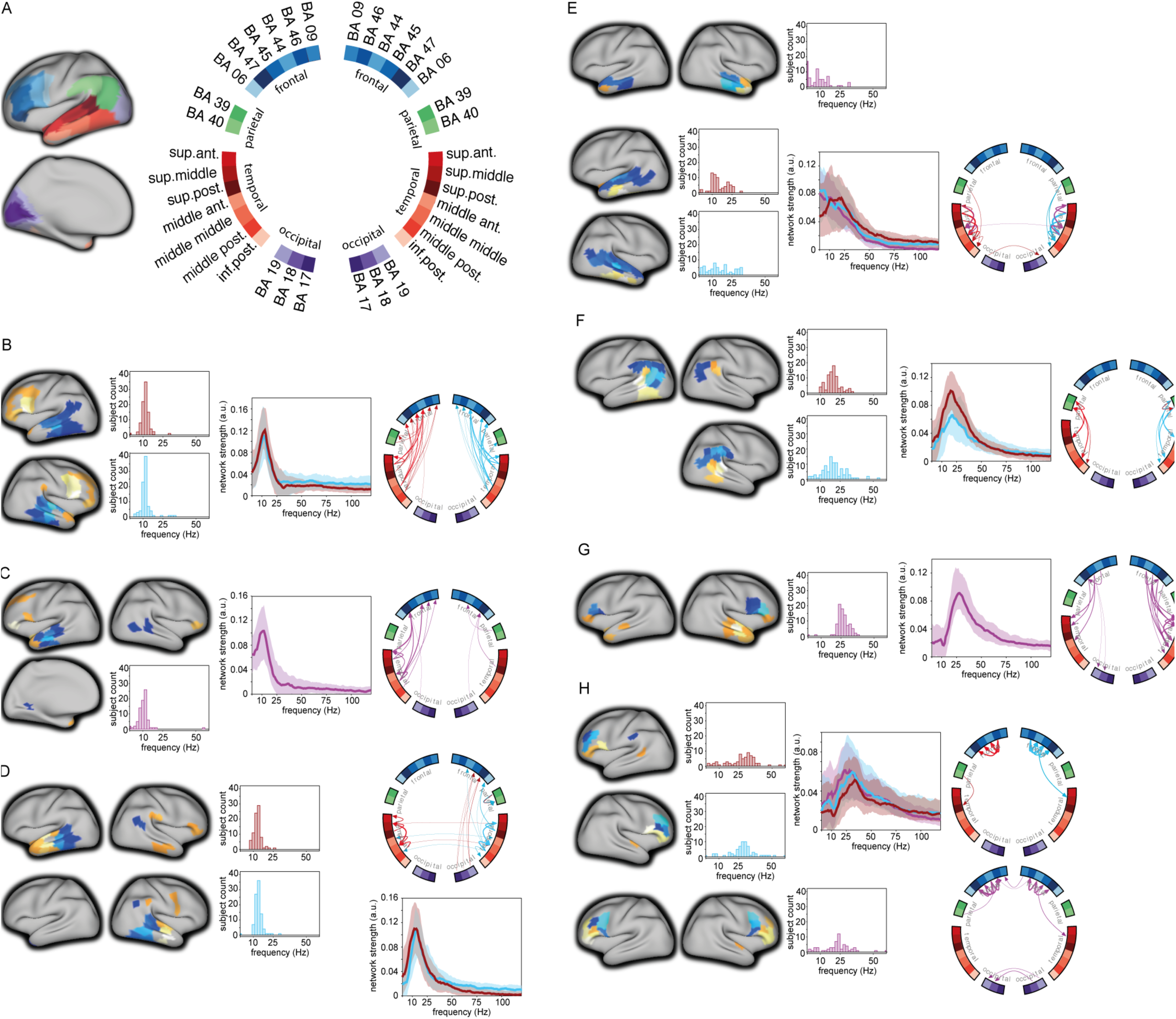
Network components obtained with non-negative matrix factorization show frequency, regional, and direction-specific interactions. (**A)**: Location of the cortical nodes, displayed on an inflated cortical sheet (left) with color coding and labelling convention (right), as used in (**B-H**). Circular grouping was according to anatomical location, using Brodmann Area (BA) labelling for the parcels outside temporal cortex, and using their relative location along the anterior/posterior and superior/inferior axis for temporal parcels). (**B-H**): Components reflecting connections between language-relevant cortical areas. Leftmost panels show the location of the parcels involved. Dark/light blue colors: regions for outflow (the lighter the color of the parcel, the stronger the relative contribution of the parcel to the component). Orange/yellow/white colors: regions for inflow. The histograms show for each of the components the distribution of the subject-specific peak frequency. The spectra show, the median (and interquartile range) spectral profiles across. The circular plots show the directed connections between the parcels. The thickness of the arrows reflects the relative strength of the connection. Components with predominantly left-hemispheric, right-hemispheric, or bilateral connections are displayed in red, light blue, and purple, respectively. (**B**): Left and right hemispheric components from temporal regions to ipsilateral frontal regions. (**C**): Component with bilateral intrahemispheric temporal to frontal connections. (**D**): Left and right hemispheric components with predominant connections from middle to anterior temporal regions. (**E**): Components with predominantly intra-temporal connections from superior to inferior regions (upper two rows), and from mid-anterior regions to the temporal pole (bottom row). (**F**): Components with predominant connections from the angular gyrus (BA 39) and supramarginal gyrus (BA 40) to posterior temporal cortex. (**G**): Component from frontal regions to temporal regions. (**H**): Components with predominantly fronto-frontal connections.

In contrast to the network components with the outflow regions in temporal cortex, the rhythmic interactions with predominant outflow from parietal (Fig. 2F) and frontal (Fig 2G-H) regions consistently showed a higher peak frequency of interaction. Components reflecting parietal to posterior temporal interactions had a median peak frequency of 20 Hz (with an IQR of 17-22 Hz and 15-26 Hz for the left and right hemispheric components, respectively), and frontal to temporal rhythmic interactions had a median peak frequency of 27 Hz (with an IQR of 25-30 Hz). Intrafrontal interactions had a somewhat more broadband spectral profile, with a median peak frequency of 24 Hz (IQR: 19-29 Hz) for directed interactions from BA 44 to BA 45/46/47. Interactions from BA 46 to BA 44/45/47 had a median peak frequency of 30 Hz (IQR: 23-35 Hz) and 29 Hz (IQR: 25-33 Hz) for left and right hemispheric connections, respectively. We statistically evaluated the peak frequency of the rhythmic interactions between components with predominant connections between parietal, frontal, and temporal brain areas (Fig. 3A). Overall, the component-specific median peak frequencies ranged from the upper end of the alpha range (12 Hz), to the upper end of the beta range (30 Hz). Moreover, components with rhythmic Granger causal outflow predominantly from temporal areas had a consistently lower peak frequency than components with Granger causal outflow from parietal or frontal areas (p<0.05, non-parametric permutation test, multiple comparison corrected). Notably, based on the NMF we could distinguish temporo-frontal interactions, with a peak frequency of 12 Hz (Fig. 2A-B, 3B-C, connection in dark red) from fronto-temporal interactions, with a peak frequency of 27 Hz (Fig. 2F, 3B-C, connection in dark blue). Figure 3B shows a schematic summary of the dominant rhythmic interactions, with the corresponding spectral profile in Fig. 3C.

**Fig 3.**
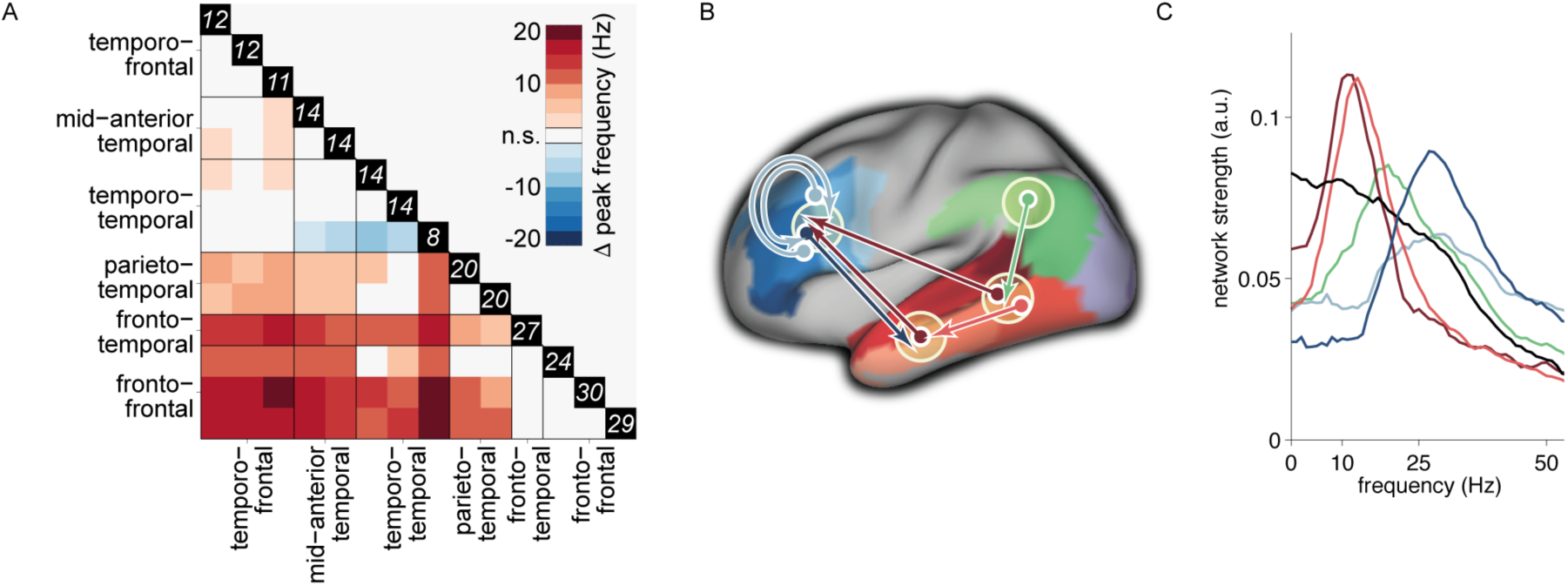
Rhythmic interactions originating from temporal/parietal cortex have consistently lower peak frequencies than those originating from frontal cortex (**A**): Pair-wise comparison between components of the peak frequencies of their corresponding spectral profiles (non-parametric permutation test, corrected for multiple comparisons). Colors represent the median across subjects of the difference in peak frequency. The values in the black boxes along the main diagonal reflect the median peak frequency for each component. (**B**): Schematic representation of the directed rhythmic cortico-cortical interactions in the language system, grouped according to the cortical output area. The temporal lobe is split into two ‘nodes’, to be able to display the rhythmic mid to anterior connection. The colored arrows refer to the spectra shown in (**C**). The black spectrum in (**C**) is the average of the components shown in (**Fig. 2E**), with dominant connections from superior temporal to middle temporal gyrus, and is not displayed as a separate connection in (**B**).

We proceeded to test whether the strength of the rhythmic interactions were modulated by the functional requirements imposed by the perceptual input. To this end, we divided the stimulus material into four conditions, based on whether the subjects were reading a well-structured sentence or a pseudo-random sequence of words (sentences and word lists), and based on the ordinal position of the words (early and late words). Importantly, we stratified the data for lexical frequency and overall signal variance, to avoid as much as possible interpretational confounds for the estimated connectivity (12, 14) due to differences univariate signal and stimulus properties (8). Subsequently we computed the Granger causal interactions for each subject and condition for the most prominent functional connections, which were extracted from the NMF-results by means of spatial clustering. We constrained the analysis to band-limited estimates of Granger causality, where the connection-specific frequency bands were obtained from the components’ peak frequencies and interquartile ranges. Contrasting sentences with sequences, we observed the strength of the interactions to be modulated from left middle temporal regions to the left temporal pole, where sequences elicited stronger interactions than sentences, and from right striate to extrastriate visual regions (Fig 4A, p<0.05, non-parametric permutation test, Holm-Bonferroni correction for multiple comparisons). Comparing early words with late words in the sentence condition showed several significantly modulated connections, with rhythmic interactions being stronger early in the sentence (Fig 4B). These connections were bilateral from temporal to frontal regions, and from middle temporal regions to the temporal pole. In addition, in the right hemisphere, we identified significantly modulated connections from frontal regions to temporal regions, and from the superior temporal gyrus to the middle temporal gyrus (p<0.05, non-parametric permutation test, Holm-Bonferroni correction for multiple comparisons). Moreover, we identified two right hemispheric connections that showed a significant interaction effect between early versus late words and sentences versus sequences (Fig. 4B).

**Fig. 4:**
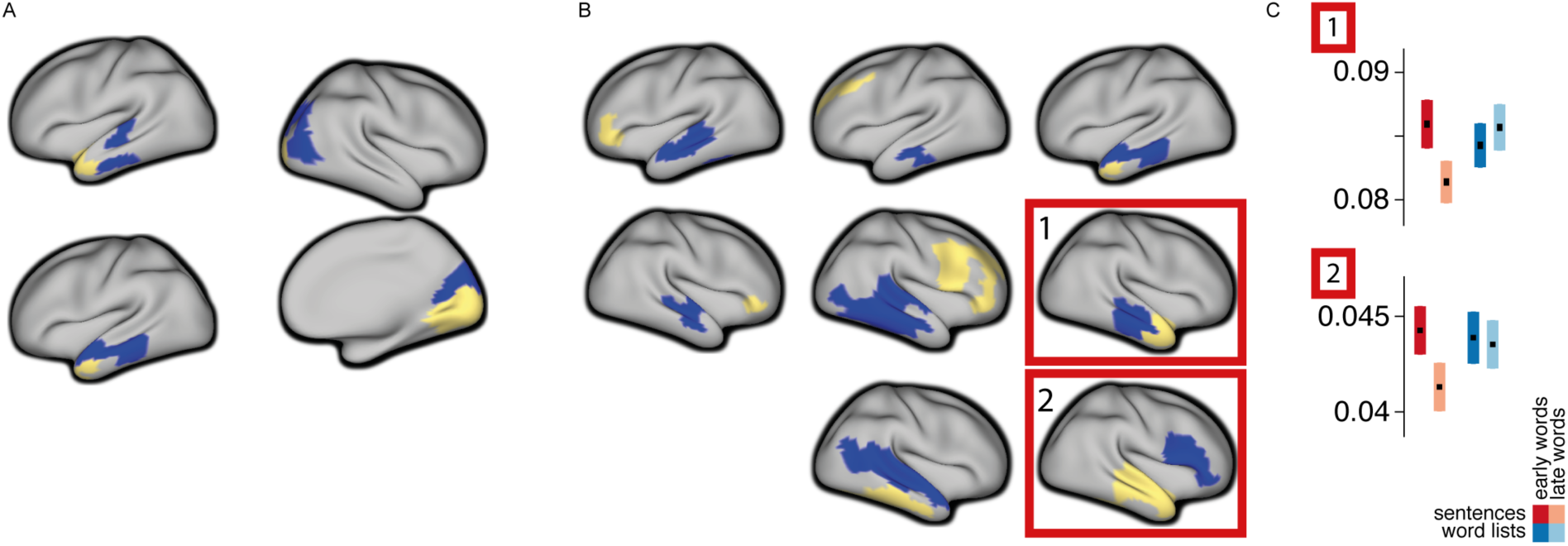
The strength of directed interactions is modulated by the linguistic input. (**A**): Connections showing stronger interactions in the word list condition than in the sentence condition (p<0.05, corrected Bonferroni-Holm). Blue parcels indicate nodes for outflow, yellow parcels indicate nodes for inflow. (**B**): Connections showing stronger interactions for early words in the sentences, compared to late words in the sentences. The connections shown in the red boxes show a significant interaction effect (p<0.05, corrected), on top of a significant early versus late effect. (**C**): Bar graphs show condition-specific mean (+/- SEM) Granger causal strength for the outlined connections.

In sum, we have provided evidence for directed interactions between cortical regions in the human brain network for language during sentence reading. Topological analysis of the overall network revealed a high degree of Granger causal inflow into anterior inferior frontal cortical regions, right anterior temporal cortex and the temporal pole bilaterally. This is in line with these regions being ‘high order’ regions, involved in the processing of more abstract features of the linguistic input, which requires integration of converging information. Frontal regions are engaged in unification operations (15), integrating lexical units into the larger context. Anterior temporal cortex is associated with conceptual object representations (16, 17).

Middle temporal cortical regions on the other hand displayed a high degree of Granger causal outflow. This is in agreement with the middle temporal gyrus’ crucial role in language comprehension at the level of single words (18, 19). Its functional connections to more anterior temporal areas, as well as to inferior frontal cortex reflect the necessity to propagate information about individual lexical items to areas that subserve integration operations. Notably, we did not observe a clear lateralization in the pattern of connections, which lends support to the evolving notion that both cerebral hemispheres are involved in the processing of linguistic stimuli (20).

Data-driven decomposition of the overall network into smaller subnetworks revealed several spatially-constrained components, corresponding with local and long range directed interactions. The clear frequency-resolved profile displayed by some of these components displayed is indicative of the interactions being mediated by rhythmic interareal synchronization. Connections originating from temporal cortical areas showed a consistently lower peak frequency (alpha, low beta) than connections originating from parietal or frontal regions (high beta). As a specific example, temporal to frontal interactions are subserved by rhythmic synchronization at ~12 Hz, whereas interactions in the opposite direction, from frontal to temporal regions peak at a frequency of ~27 Hz.

At first glance, these findings correspond well with recent work in the visual system. There it was shown that feedforward and feedback connections, as defined by their characteristic cortical laminar connectivity profile (21), could be distinguished in terms of their frequency of interaction (7, 22, 23). However, in the visual system, feedforward connections have been functionally characterized by gamma band synchronization (>50 Hz) (and to a lesser extent by theta band synchronization) and feedback connections by alpha/beta synchronization. A Granger causality spectral peak in the gamma frequency range was absent in our data. While studies in the visual system allow for experimental paradigms eliciting robust gamma band rhythmicity, language paradigms do not typically lead to very distinct local gamma band responses (24), rendering the likelihood of detecting gamma band interactions low. In addition, characterization of feedforward and feedback connections based on their cytoarchitectonic connectivity profiles is likely to be more distinct in peripheral sensory systems than in higher cortical regions (25), such as the higher order areas in the human brain network for language. Consequently, there is no reason to assume that the emergent functional properties of the language network, in terms of the frequency of interactions should directly map onto observations in the visual system. Nevertheless, our data reveal frequency-specific subnetworks in the brain system for language.

Further exploration of the potential functional significance of these interactions revealed that the linguistic context modulates left lateralized mid-temporal to anterior temporal interactions, as well as right lateralized extrastriate to striate interactions. In a sentential context, the incremental availability of contextual information allows for the generation of constraining predictions about the upcoming input, likely facilitating processes such as lexical selection. Bilateral interactions from temporal to frontal regions, from mid-temporal to anterior temporal regions, from right lateralized frontal to temporal regions, and from superior temporal to middle temporal regions are stronger early in the sentence, when the constraining context is still relatively weak, as opposed to later in the sentence. These stronger interactions early in the sentence might reflect the increased need for information exchange between these regions, in order to establish a linguistic context.

In conclusion, this study shows directional interactions in the highly dynamic cortical network of language-relevant areas, with salient differences in the specific frequencies that support the communication protocols in the temporo-frontal and the fronto-temporal directions. Although our findings are in line with earlier reports of a frequency difference between feedforward and feedback connections, the carrier frequencies in the language network shown here deviate from what has been observed in the visual system. Yet, the effect of linguistic context on the strength of some of these connections suggests the functional relevance of dynamic rhythmic cortical interactions during cognitive processing in general, and language processing in particular (26).

## Acknowledgments

This work was funded by the Royal Netherlands Academy of Sciences (Spinoza award to P.H.) and the Dutch Organization for Scientific Research (NWO gravitation grant to P.H., and an NWO VIDI grant to J.M.S.).

## Materials and Methods

### Experimental procedure and MEG data acquisition

A total of 102 native Dutch speakers (51 males), with an age range of 18 to 33 years (mean of 22 years), participated in the experiment. All participants were right-handed, had normal or corrected-to-normal vision, and reported no history of neurological, developmental or language deficits. The study was approved by the local ethics committee and followed the guidelines of the Helsinki declaration. Participants received monetary compensation for their participation.

The participants were seated comfortably in a magnetically shielded room, and were instructed to read sequences of words (total number of 240, with 9 to 15 words per sequence), which were presented sequentially on a backprojection screen, placed in front of them. All words were presented at the center of the screen within a visual angle of 4 degrees, in a black mono-spaced font, on a grey background using Presentation software (Version 16.0, Neurobehavioral Systems, Inc). The vertical refresh rate of the LCD projector was 60 Hz. The sequences of words formed either well-formed sentences, or consisted of a scrambled version of a sentence, where the word order was randomly shuffled. For the remainder we refer to these latter stimulus sequences as word lists. See Lam et al. 2016 (24) for more details about the stimulus material used. Sentences and word lists were presented in small blocks, of 5 sentences (or word lists) each, to a total of 120 stimuli per condition. In order to check for task compliance, in a random 10% of the word sequences, they were followed by a yes/no question about the content of the previous sentence/word list.

MEG data were collected with a 275 axial gradiometer system (CTF). The signals were analog lowpass filtered at 300 Hz, and digitized at a sampling frequency of 1200 Hz. The participant’s head was registered to the MEG-sensor array using three coils, attached to the participant’s head (nasion, left and right ear canals). Throughout the measurement the head position was continuously monitored using custom software (27). During breaks the participant was allowed to reposition to the original position if needed. Participants were able to maintain a head position within 5 mm of their original position. Three bipolar Ag/AgCl electrode pairs were used to measure the horizontal and vertical electro-oculogram, and the electro-cardiogram.

### Artifact rejection and subtraction of single-trial activity

All analyses were done with custom written Matlab scripts and Fieldtrip (28). Data were initially epoched from −100 to 600 milliseconds relative to word onset. Segments contaminated by artifacts due to eye movements, muscular activity and SQUID jumps were discarded prior to further analysis. Next, we subtracted the event-related response from the single trial data with the ASEO algorithm (29). The acronym ASEO stands for Analysis of Single-trial ERP and Ongoing activity, and the aim of the application of this algorithm in this context was to attenuate the effects of evoked transients on the estimation (and subsequent interpretation) of Granger causality (30). Transients in the signals violate the underlying assumption of stationarity, and moreover may result in non-zero Granger causality estimates, due to systematic latency differences of the peak of the transient signals across regions. Although such latency differences may reflect an actual interaction (where temporal precedence of a transient signal peak in region A, compared to region B, may be an indication that A is causing B), their spurious effect on the estimated Granger causality is unwanted, if the aim is to interpret the frequency domain Granger causality in terms of directed synchronized interactions. In our experimental setup, transient brain responses could not be avoided (as opposed to for instance (7)), and we developed a procedure (performed at the sensor-level) to attenuate the effect of transient evoked components, combining the ASEO algorithm with a blind source separation technique (denoising source separation (DSS) (31)). In short, the ASEO algorithm models single-trial signals as a combination of ongoing activity and event-related components, where the latter are modelled as a set of ‘canonical’ components, each with a trial-specific latency and amplitude. In a typical application (32), the single-trial estimates of latencies and amplitude are used as dependent variables for subsequent analysis. Here, however, we subtracted the single-trial evoked responses that were reconstructed from the latency and amplitude estimates, which results in a better account of ongoing activity, as compared to the subtraction of a fixed average event-related response from each signal. DSS was used to iteratively unmix the sensor-level data into a set of components, where the DSS framework allows for the unmixing algorithm to capitalize on specific features of the requested components. Specifically, we applied an iterative procedure, where each iteration consisted of the following steps:

1. Estimation of the dominant DSS-component using quasi-periodic averaging, which essentially extracts components with strong evoked transients, time locked to word onset. This step yields a spatial map of mixing weights, describing for each MEG sensor the extent to which this component is present in the MEG signals, as well as an observations-by-time matrix of the component time series.
2. Application of the ASEO-algorithm to the single-trial time series of the estimated DSS-component, yielding single trial estimates of the evoked transients.
3. Backprojection of the component’s single-trial evoked transients to the MEG sensor-level, using the spatial map of mixing weights, obtained in step 1.
4. Subtraction of the backprojected evoked transients from the MEG sensor-data, yielding MEG-sensor data that served as input data for the next iteration.

We performed 5 iterations, i.e. we removed 5 timelocked components. Removal of additional components did not affect the global field power appreciably (Supp.fig.1E). The different steps and the effect of this cleaning procedure are illustrated in Supplementary Figure 1.

### Source reconstruction and parcellation of source-reconstructed activity

After the cleaning of the sensor data with the combined DSS-ASEO procedure, we performed source reconstruction using a linearly constrained minimum variance beamformer (LCMV) (33). For this, we computed the covariance matrix between all MEG-sensor pairs, as the average covariance matrix across the cleaned single trial covariance estimates. This covariance matrix was used in combination with the forward model, defined on a set of 8196 locations on the participant-specific reconstruction of the cortical sheet to generate a set of spatial filters, one filter per dipole location. Individual cortical sheets were generated with the Freesurfer package (version 5.1) (http://surfer.nmr.mgh.harvard.edu), coregistered to a template based on a surface-based coregistration approach, with a dedicated script using the Caret software (http://brainvis.wustl.edu/wiki/index.php/Caret:Download, http://brainvis.wustl.edu/wiki/index.php/Caret:Operations/Freesurfer_to_fs_LR), and subsequently downsampled to 8196 nodes, using the MNE software (http://martinos.org/mne/). The forward model was computed using Fieldtrip’s ‘singleshell’ method (34), where the required brain/skull boundary was obtained from the subject-specific T1-weighted anatomical images. Next, we applied an atlas-based parcellation scheme to further reduce the dimensionality of the data. To this end, we used the Conte69 atlas (http://brainvis.wustl.edu/wiki/index.php//Caret:Atlases/Conte69_Atlas), which provides a parcellation of the neocortical surface, based on Brodmann’s cytoarchitectonic atlas, consisting of 41 labelled parcels per hemisphere. This parcellation scheme was further refined, breaking up the larger parcels into a set of subparcels, respecting the original boundaries (e.g.: breaking up the middle temporal gyrus in smaller parcels along the anterior/posterior axis). This resulted in a parcellation scheme consisting of 191 parcels per hemisphere.

For each parcel we obtained a parcel-specific spatial filter as follows: we concatenated the spatial filters of the vertices comprising the parcel, obtained a set of time courses of the event-related field at each parcel, and performed a principal component analysis on the result. We selected for each parcel the first 2 spatial components explaining most of the variance in the signal. For the parcels used, the two dominant spatial components explained on average 90% of the signal variance within each parcel (range: 74%-96%).

### Preselection of the connections between language-relevant areas

For the connectivity analysis, we constrained ourselves a priori to a subset of connections between parcel-pairs, using known ‘long range’ macro-anatomical fibre pathways between parcels comprised of core language regions and the visual system as described in the literature (1, 9–11). This preselection was motivated by the fact that direct functional connections should be supported by direct anatomical connections. In addition, we allowed a priori for direct connections between neighboring nodes, which is a fair assumption given the characteristics of cortico-cortical connections observed in anatomical tracing studies (e.g. (25, 35)), where local connections are abundant. We included intrahemispheric connections from both hemispheres, and also included interhemispheric connections between homologous areas. The nodes were defined based on the labelling scheme of Brodmann, where each of these nodes could consist of one or more subparcels (where subparcels were defined as described above). In addition, nodes in temporal cortex were classified according to their position along the anterior-posterior axis (distinguishing anterior, middle and posterior parts), and along the superior-inferior axis (distinguishing superior, middle and inferior parts). Figure 2A in the main text shows how the individual nodes were labeled. As major long-range fiber pathways we included the arcuate fasciculus (AF), the superior longitudinal fasciculus (SLF), the extreme capsule (EC), the uncinate fasciculus (UC), and the inferior fronto-occipital fasciculus (IFOF). The AF provides widespread connections between the temporal cortex (predominantly the middle and superior temporal gyri) and various frontal areas (BA44/45/6/9). The SLF connects frontal areas (notably BA44) with posterior superior temporal, and parietal areas. The EC connects frontal areas with the middle part of superior and middle temporal gyrus. The UC connects frontal areas with the temporal pole, and the IFOF connects frontal areas with occipital areas. Connections between directly adjacent parcels were excluded for further analysis to reduce spurious estimates of connectivity due to spatial leakage of source reconstructed activity. The selection scheme resulted in 4350 connections between pairs of parcels, which notably consisted of a sparse subset of all possible pairwise connections between the 156 parcels used for the Granger causality analysis.

### Granger causality computation and statistical evaluation of overall network topology

For computational efficiency, we computed the spectral representation of the signals at the sensor-level, and projected this into source space, using the parcel-specific spatial filters. The spectral representation of the signals was obtained using the Fast Fourier transform in combination with multitapers (using 5 Hz smoothing), on the time domain data from 200 until 600 ms after word onset. The sensor-level Fourier transformed data was projected into source space, and for each pair of parcels we computed the cross-spectral density matrix. Subsequently we performed non-parametric spectral matrix factorization for each pair of parcels, followed by computation of Granger causality (36, 37). We computed Granger causality based on the source-projected Fourier transform of time-reversed data, where time-reversal is essentially equivalent to complex conjugation of the Fourier coefficients, in order to distinguish ‘weak’ asymmetries from ‘strong’ asymmetries, as described by Haufe et al. (13, 38). Essentially, a weak asymmetry is an apparent directional interaction between a pair of network nodes, which is the consequence of a difference in signal-to-noise ratio across nodes (14), and difficult to avoid when the signals consist of a linear mixture of underlying sources (39). We compared Granger causality with reverse Granger causality, and selected only parcel pairs for subsequent analysis for which the parametric null-hypothesis of the means (across subjects) could be rejected at a p-value < 0.05, corrected for multiple comparisons (one sided T-test, with Bonferroni correction). This reduced the number of connections that were used for subsequent analysis from 4350 to 713. Next, we evaluated the topology of this resulting network by quantifying the node degree for each of the 156 parcels involved, identifying ‘hubs’ for inflow and outflow (figure 1). We quantified the probability of observing the computed node degree under the null hypothesis of the 713 connections being a random subset of the originally included 4350 connections, using a permutation test. Using Bonferroni correction, a p-value of 0.05/(2*156)=1.6x10^-4^ was considered significant (each of the 156 parcels was tested twice, once for the degree for inflow, and once for the degree for outflow). As an important control analysis, we computed, across parcels, the Spearman’s rank correlation between the inflow and outflow degree on the one hand, and signal variance, the norm of the spatial filter, and the signal-to-noise ratio (SNR) on the other hand. The norm of the spatial filter corresponds with an estimate of the projected noise, and the ratio between the signal variance and the spatial filter’s norm corresponds with an SNR estimate. The motivation for this analysis is to check whether there is a relationship between the node degree and simple univariate signal(-to-noise) properties, which may give rise to spurious inferences about the directionality of estimated interactions (14). Specifically, assuming the worst, one could hypothesize that parcels with a large degree of inflow (outflow) also show on average a low (high) signal(-to-noise), when comparing across parcels. The results of this control analysis are shown in Supplementary Table 1. Based on this analysis, which did not reveal any significant correlations, we argue that the observed patterns of node degree in the brain network for language are not consequences of systematic differences in univariate signal properties.

### Non-negative matrix factorization and network visualization

We explored the network topology by performing non-negative matrix factorization (NMF) with sparsity constraints (40) on the resulting Granger causality spectra. The purpose of this analysis is to describe the reconstructed connectivity data as a low-dimensional mixture of network components, each of which with a subject-specific spectral profile. We opted for NMF, because the non-negativity constraint facilitates the interpretation of the components (as opposed to e.g. a statistical independence constraint as applied in independent component analysis). This is because Granger causality is strictly non-negative. The data matrix that was subjected to the factorization algorithm was constructed by concatenating across subjects Granger causality spectra, normalized for the standard deviation per subject. The columns in this matrix reflect the individual connections (across subjects and frequencies), and the rows in this matrix (number of frequency bins times number of subjects) reflected the connections for a given frequency bin and subject. Typically, the outcome of NMF is dependent on the number of components (which has to be chosen a priori), and of the initial random starting conditions. We explored a range of ‘number of components’ but settled on the number 20 for the remainder of the paper, because this number provided a reasonable balance between providing a small number of easily interpretable components, while at the same time maintaining a good separation between subnetworks. We used the ‘icasso’ framework (41), which applies a hierarchical clustering procedure on the outcome of repeated decompositions (which differ due to the random starting conditions). For this we used 40 repeated random initializations to estimate the components that robustly represent the underlying structure of the data, irrespective of the random initializations of the NMF algorithm.

The outcome of this procedure consisted of two matrices. One matrix represents the network component spatial fingerprints, quantifying for each of the edges its relative contribution to the network components. The other matrix contains for each network component a subject-specific spectral profile, quantifying the subject-wise relative and frequency-specific contribution to the network components. For visualization purposes, we assigned each of the edges to a unique network component based on their relative weight. Subsequently, the different aspects of the network components were depicted as follows: To obtain spatial maps of the nodes (i.e. anatomical parcels) participating in a particular component, we summed across each node’s contributing edges the outflow and inflow separately, and displayed these onto an inflated representation of the cortical sheet, using hcp-workbench (http://www.humanconnectome.org/software/connectome-workbench.html

To visualize the connections between the parcels, for each pair of parcels we averaged the connection weights across all pairs of subparcels that constituted the parcel pair. Connections where drawn as directed arrows, where the thickness of the lines reflects the overall weight of the connection. The spectral profiles of the network components were visualized as the median across subjects and their interquartile range. For display purposes we ordered the components according to their dominant regions for outflow. Figure 2 in the main text shows the components that are made up predominantly by connections between language-relevant cortical parcels.

Components that are made up predominantly by connections between visual cortical parcels, as well as components with spatially very diffuse connections are displayed in supplementary figure 2.

### Condition-specific statistical evaluation

To investigate whether the involvement of the network components was modulated by functional constraints of the linguistic input, we estimated condition-specific Granger causality in the dominant connections extracted from the identified network components. The individual conditions were defined according to whether the words were presented in a well-formed sentence context (or were part of a word list), and according to whether the words were presented early in the sentence/word list (words 2-4), or late in the sentence/word list (n-3 until n-1, with n the number of words in the sentence/word list). In order to account for potential interpretational confounds of the resulting Granger causality estimates we adopted a stratification procedure to ensure that, for each of the parcel pairs in each of the subjects, the marginal distributions of the epoch-wise signal variances as well as the words’ lexical frequencies were equalized across conditions. Lexical frequencies were estimated using the Subtlex-NL database (http://crr.ugent.be/programs-data/subtitle-frequencies/subtlex-nl). Condition-specific histograms for lexical frequency were generated using 13 log-spaced bins. Histograms for signal variance were generated using 6 log-spaced bins. The consequence of this procedure is that only a subset of epochs is used for the subsequent estimation of Granger causality, where the parcel-pair specific number of epochs varies across parcel pairs. On average 50% of the epochs were retained (range: 20-75%), corresponding to 147 (range: 45-235) epochs.

From each of the extracted network components we defined a dominant connection as a spatially clustered set of edges that fulfilled the following criteria:

- Each cluster consisted of at least 4 edges.
- The inflow/outflow nodes consisted of spatially adjacent cortical parcels.
- Nodes that for a given cluster of edges served both as input and output node were discarded, as well as the edges to which these nodes contributed.

This resulted in 42 connections for which we computed subject and condition specific Granger causality, as an average across the contributing edges, and across the component specific frequency range, defined by the interquartile range across subjects. We performed a non-parametric permutation test to evaluate the following contrasts:

1. Sentence – word list words
2. For the sentence condition: early – late words
3. Interaction effect: (early-late words sentences) – (early-late words sequences).

The statistical test performed was a two-sided permutation test (using 20,000 permutations) on Wilcoxon’s signed rank statistic (Z-score) with a Bonferroni-Holm stepdown control for the family-wise error rate.

---

**Fig. S1.**
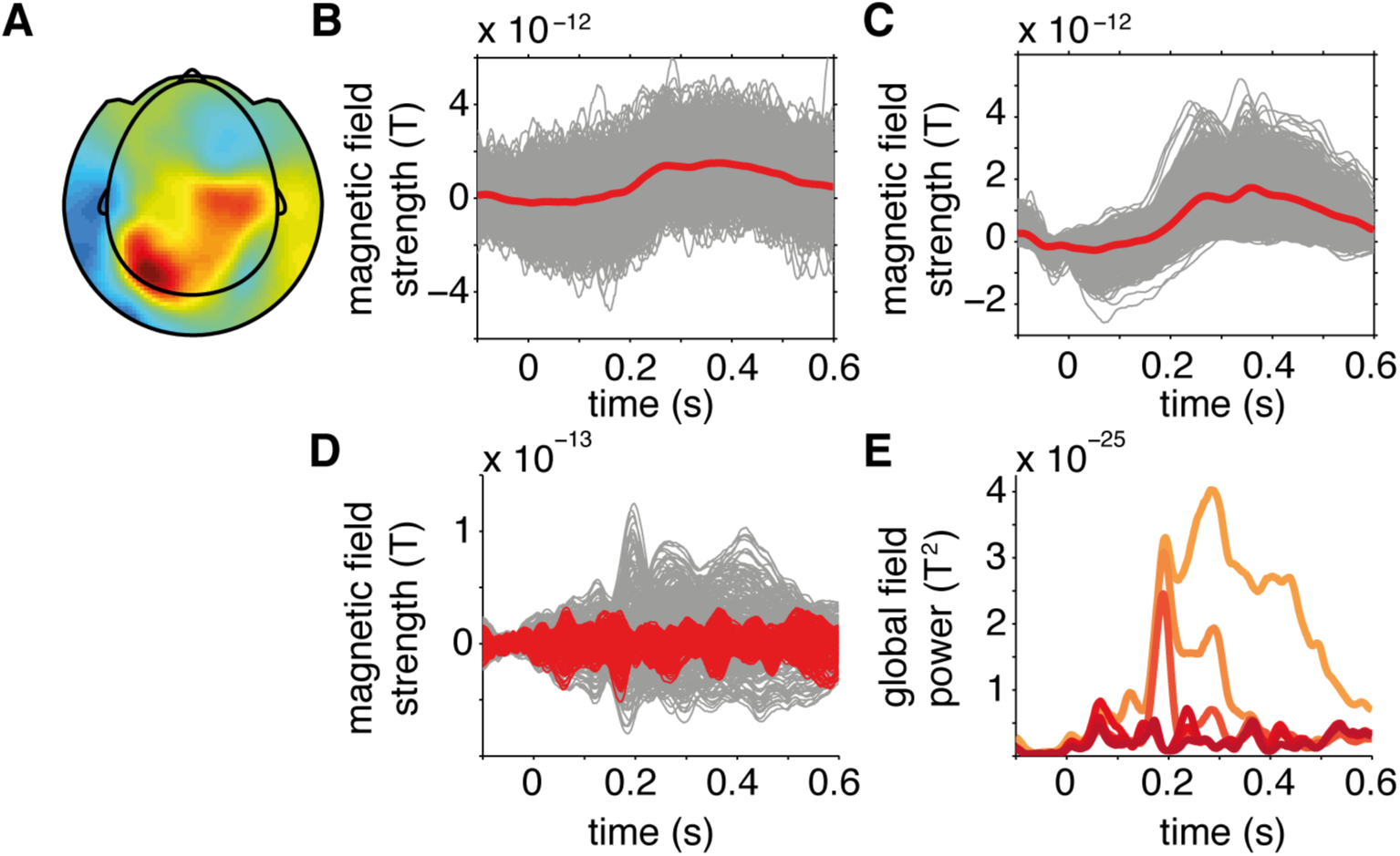
Illustration of combined DSS/ASEO procedure tor the removal of word-onset event-related signal transients. (A): spatial topography of the mixing coefficients for the first extracted DSS-component, for an example subject. (B): single trial time courses (in gray) of the first DSS-component, time-locked to word onset, average across trials in red. (C): single trial estimates of the stimulus-locked transient response, estimated with the ASEO algorithm, average across trials in red. (D): overlay of single channel event-related averages before (gray) and after (red) the cleaning procedure. (E): global field power across channels of the event-related average (light orange) and after iterative removal of 5 DSS components (colors going from orange to dark red).

**Fig. S2.**
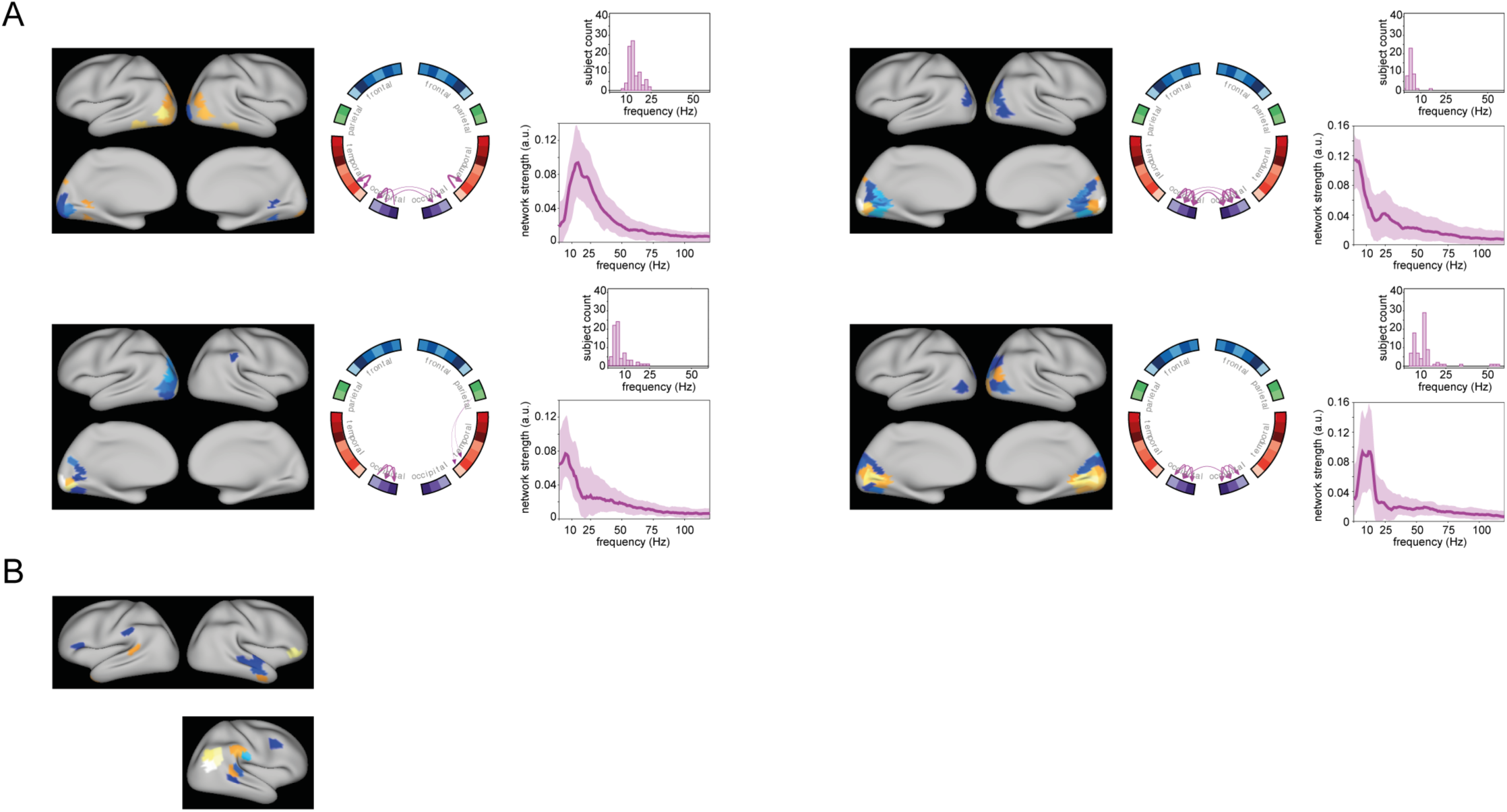
Network components obtained by group-level non-negative matrix factorization with predominant connections between visual cortical areas (A), and with spatially diffuse connections (B).

**Table S1.** Correlation between node degree and univariate signal properties.

|  | *Node degree inflow* |  | *Node degree outflow* |  |
| --- | --- | --- | --- | --- |
|  | *Correlation* | *P-value* | *Correlation* | *P-value* |
| *Signal variance* | 0.054 | 0.51 | 0.077 | 0.34 |
| *Spatial filter norm* | 0.096 | 0.23 | 0.12 | 0.15 |
| *SNR* | -0.097 | 0.23 | 0.032 | 0.69 |

